# Off-target depletion of plasma tryptophan by allosteric inhibitors of BCKDK

**DOI:** 10.1101/2024.03.05.582974

**Authors:** Caitlyn E. Bowman, Michael D. Neinast, Cholsoon Jang, Jiten Patel, Megan C. Blair, Emily T. Mirek, William O. Jonsson, Qingwei Chu, Lauren Merlo, Laura Mandik-Nayak, Tracy G. Anthony, Joshua D. Rabinowitz, Zolt Arany

## Abstract

The activation of branched chain amino acid (BCAA) catabolism has garnered interest as a potential therapeutic approach to improve insulin sensitivity, enhance recovery from heart failure, and blunt tumor growth. Evidence for this interest relies in part on BT2, a small molecule that promotes BCAA oxidation and is protective in mouse models of these pathologies. BT2 and other analogs allosterically inhibit branched chain ketoacid dehydrogenase kinase (BCKDK) to promote BCAA oxidation, which is presumed to underlie the salutary effects of BT2. Potential “off-target” effects of BT2 have not been considered, however. We therefore tested for metabolic off-target effects of BT2 in *Bckdk*^-/-^ animals. As expected, BT2 failed to activate BCAA oxidation in these animals. Surprisingly, however, BT2 strongly reduced plasma tryptophan levels and promoted catabolism of tryptophan to kynurenine in both control and *Bckdk*^-/-^ mice. Mechanistic studies revealed that none of the principal tryptophan catabolic or kynurenine-producing/consuming enzymes (TDO, IDO1, IDO2, or KATs) were required for BT2-mediated lowering of plasma tryptophan. Instead, using equilibrium dialysis assays and mice lacking albumin, we show that BT2 avidly binds plasma albumin and displaces tryptophan, releasing it for catabolism. These data confirm that BT2 activates BCAA oxidation via inhibition of BCKDK but also reveal a robust off-target effect on tryptophan metabolism via displacement from serum albumin. The data highlight a potential confounding effect for pharmaceutical compounds that compete for binding with albumin-bound tryptophan.

## Introduction

The essential branched chain amino acids (BCAAs: leucine, valine, and isoleucine) comprise up to a third of the protein content in muscle and can be an important energy source in catabolic muscles ^1^. Interest in BCAAs grew after unbiased metabolomic profiling in large retrospective and prospective studies, as well as genetic studies, showed that elevated serum BCAAs correlate with insulin resistance (IR) ^2-6^. A Mendelian randomization study also showed that polymorphisms that cause elevations in circulating BCAAs also predict IR ^7^. Similarly, genetic associations with insulin resistance have been causally implicated with elevated BCAAs, suggesting a potential positive feedback loop ^8,9^. Elevations in circulating BCAAs have also been associated with cardiovascular disease ^10-14^ and pancreatic cancer^15,16^.

These clinical associations have inspired mechanistic studies in rodent models. Nearly all of these have relied on a small molecule, *3*,*6-dichlorobenzo(b)-thiophene-2-carboxylic acid* (BT2), which activates BCAA oxidation ^17^. Systemic administration of BT2 lowers plasma BCAA levels and improves IR in multiple contexts ^18-21^. Sodium phenylbutyrate (NaPB), a BT2-like molecule, was recently shown in a small, randomized, placebo-controlled, double-blinded clinical trial to promote insulin sensitivity in diabetic subjects ^22^. Activation of BCAA catabolism by BT2 is protective in various models of heart failure, including myocardial infarction, transverse aortic constriction, and ischemia/reperfusion ^23-25^, and in various tumor models^26-28^.

BT2 promotes BCAA catabolism by allosterically inhibiting the activity of branched chain ketoacid dehydrogenase kinase (BCKDK) ^17,29^ (Figure 1A). BCKDK phosphorylates and inhibits the branched chain ketoacid dehydrogenase (BCKDH) complex, which carries out the rate-limiting step of BCAA catabolism by decarboxylation and dehydrogenation of branched chain ketoacids (BCKAs), yielding coenzyme A (CoA)-esterified intermediates that are further metabolized in the mitochondrial matrix. Inhibition of BCKDK thus promotes BCAA oxidation and lowers plasma BCAAs in rodent models. Multiple pharmaceutical companies are actively developing novel BT2-based small molecule series ^20,30,31^. BT2 and several analogs, including NaPB, bind BCKDK in the same allosteric pocket as does the leucine-derived BCKA, thus mimicking endogenous autoinhibition ^17,29^. Other BT2-protein interactions have not been reported, and BT2 is widely thought to be specific.

**Figure 1.**
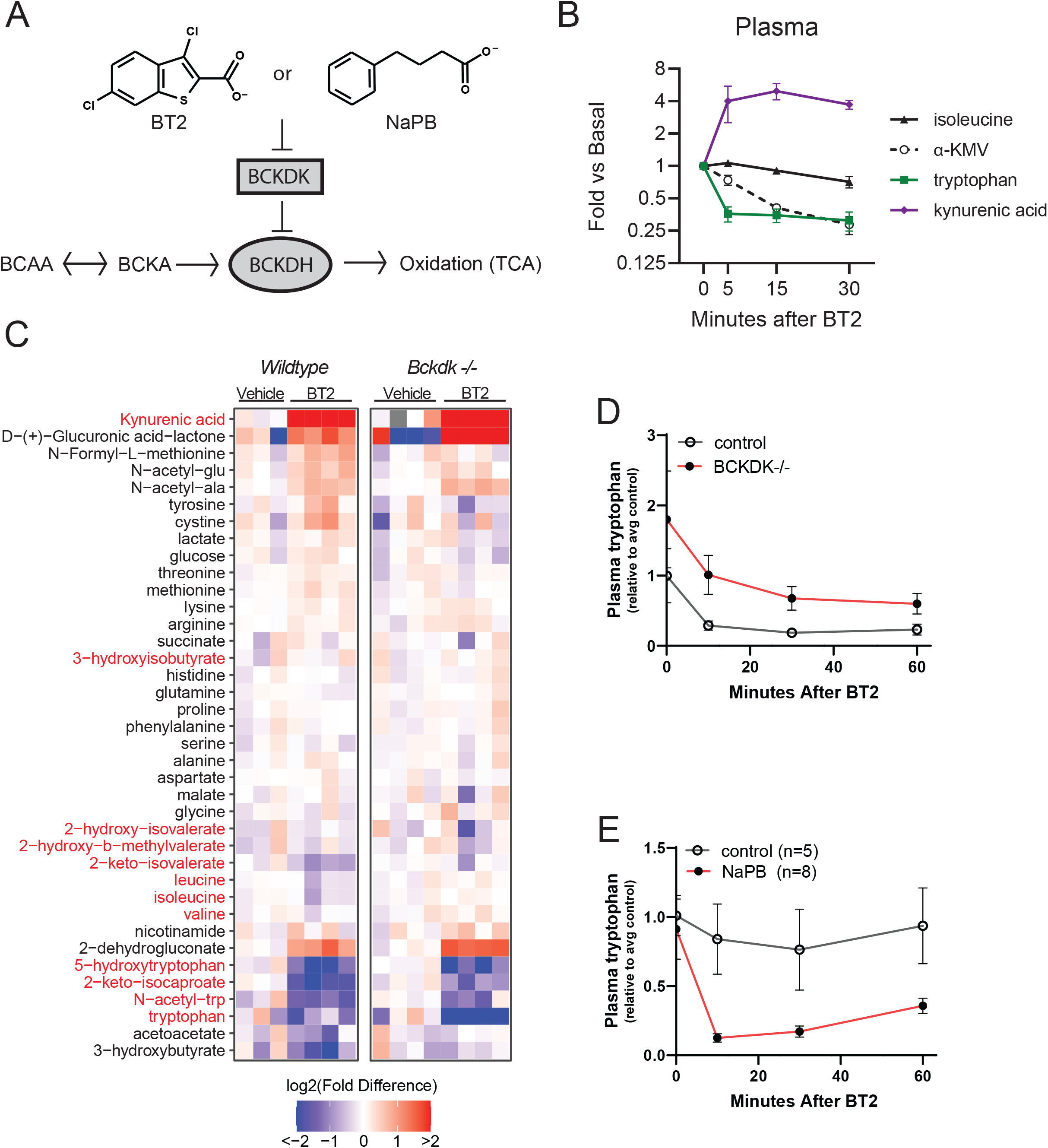
Inhibitors of BCKDK activate tryptophan catabolism in a BCKDK-independent manner. (**A**) BT2 or sodium phenylbutyrate (NaPB) inhibit BCKDK to activate BCAA oxidation. BT2 (40 mg/kg) was injected through a jugular vein catheter, and arterial plasma was sampled via a catheter in the carotid artery. Metabolite levels measured by LC-MS were normalized to the plasma sample collected before BT2 injection (n=3). Kynurenic acid was not detected in the 0-minute samples, so its value was imputed from the limit of detection in other experiments. (**C**) Whole-body BCKDK^-/-^ knockout (KO) mice were fasted 5 hours then given BT2 (40 mg/kg, i.p.) or saline, and plasma metabolites were measured after 30 minutes by LC-MS (n=4). Red highlighted metabolites are BCAA- or tryptophan-related metabolites. The fill color indicates the fold difference (log2) between the abundance of the compound in each sample and the mean of vehicle-treated samples for each genotype. (**D**) Acute oral gavage of BT2 (40 mg/kg) reduced plasma tryptophan to a similar extent in both control (n=5) and BCKDK KO mice (n=7) (time effect p=0.0278 and genotype effect p=0.0307 in mixed-effects model). (**E**) NaPB injection (200 mg/kg, i.p.) also depleted plasma tryptophan (n=5 control; n=8 NaPB). BCKDK = branched chain ketoacid dehydrogenase kinase; BCKDH = branched chain ketoacid dehydrogenase; BCAA = branched chain amino acid; BCKA = branched chain ketoacid; TCA = tricarboxylic acid cycle; α-KMV = alpha-ketomethylvalerate.

In the course of our studies with BT2, we observed that, concomitantly with reduced plasma BCKAs, tryptophan levels are also rapidly and robustly reduced in the circulation of mice treated with BT2 ^32^. This decrease in circulating levels of the essential amino acid tryptophan has also been previously reported by Zhou and colleagues in supplemental information but never discussed ^19^. The strong suppression of plasma tryptophan levels cannot easily be explained by inhibition of BCKDK. We therefore set out to identify the molecular mechanism by which BT2 enhances tryptophan catabolism.

## Results

### BT2 activates tryptophan catabolism independently of BCKDK

Acute administration of BT2 to mice rapidly and robustly decreased plasma tryptophan levels, more rapidly than plasma BCKAs (Figure 1B) ^21,32^. Metabolic fates of tryptophan include serotonin biosynthesis and catabolism via the kynurenine pathway (reviewed in ^33-35^). After BT2 administration, circulating levels of kynurenic acid increased dramatically (Figure 1B). We conclude that BT2 enhances tryptophan catabolism in one or multiple cell types.

We next sought to determine the molecular target of this BT2-induced increase in tryptophan catabolism. We first tested the presumption that BCKDK may be required for the observed suppression of circulating tryptophan. *Bckdk*^-/-^ mice and wild-type controls were injected intraperitoneally (i.p.) with saline or 40 mg/kg BT2 and plasma was collected 30 minutes post-injection (Figure 1C). Compared to control mice, *Bckdk*^-/-^ mice have reduced BCKAs at baseline, which are not further reduced by BT2 treatment, confirming that the activation of BCAA catabolism by BT2 requires BCKDK, as we previously showed ^23^. Interestingly, *Bckdk*^-/-^ mice had elevated plasma tryptophan levels at baseline compared to littermate controls (Figure 1C, D). Surprisingly, however, the plasma tryptophan levels were reduced by BT2 treatment in *Bckdk*^-/-^ mice to a similar degree as in control mice (Figure 1C). Oral gavage of BT2, which is the more common dosing method, also rapidly depleted plasma tryptophan in both *Bckdk*^-/-^ and littermate controls (Figure 1D). Treatment with NaPB, a less potent, BT2-like inhibitor of BCKDK, similarly reduced plasma tryptophan (Figure 1E). NaPB is used clinically as a treatment for uremic diseases (unrelated to BCKDK inhibition), and we previously reported reduced tryptophan levels in patients given NaPB ^22^. We conclude that BT2 promotes tryptophan catabolism by targeting a molecule other than BCKDK.

### BT2 lowers plasma tryptophan independent of predominant tryptophan catabolic pathways

The rapid and robust appearance of kynurenic acid in plasma following BT2 administration (Figure 1B) strongly suggested that BT2 may activate this tryptophan catabolic pathway directly. We thus focused our attention on the rate-limiting catabolic steps immediately downstream of tryptophan (Figure 2A). Tryptophan is first converted to kynurenine via the tryptophan 2,3-dioxygenase (TDO) family of enzymes ^33-35^, which include three members: TDO2, IDO1, and IDO2 (IDOs are named for indole 2,3-dioxygenase). The tryptophan 2,3-dioxygenase reaction is the rate-limiting step in tryptophan catabolism and occurs largely in the liver, where TDO2 is the dominant enzyme responsible ^35,36^. We therefore generated *Tdo2*^-/-^ mice by CRISPR/Cas9-mediated genome editing and deletion of *Tdo2* exons 1 and 2. *Tdo2*^-/-^ mice were born at the expected Mendelian ratios and exhibited no overt phenotypes. Plasma tryptophan levels in *Tdo2*^-/-^ mice were more than 8-fold higher than littermate controls (Figure 2B), as has been reported previously in a different murine model of Tdo2 deficiency ^37-39^. Importantly, however, the plasma tryptophan levels were reduced by BT2 treatment in *Tdo2*^-/-^ mice with the same kinetics as in control mice (Figure 2B). Interestingly, tryptophan levels rebounded faster after BT2 treatment in *Tdo2*^-/-^ mice than in control animals, perhaps due to the baseline hypertryptophanemia caused by loss of the rate-limiting enzyme in the pathway. Phospho-BCKDH levels were similarly decreased in the livers of BT2-treated mice of both genotypes (Figure 2C). We conclude that BT2 promotes tryptophan catabolism by targeting a molecule other than the rate-limiting TDO enzyme.

**Figure 2.**
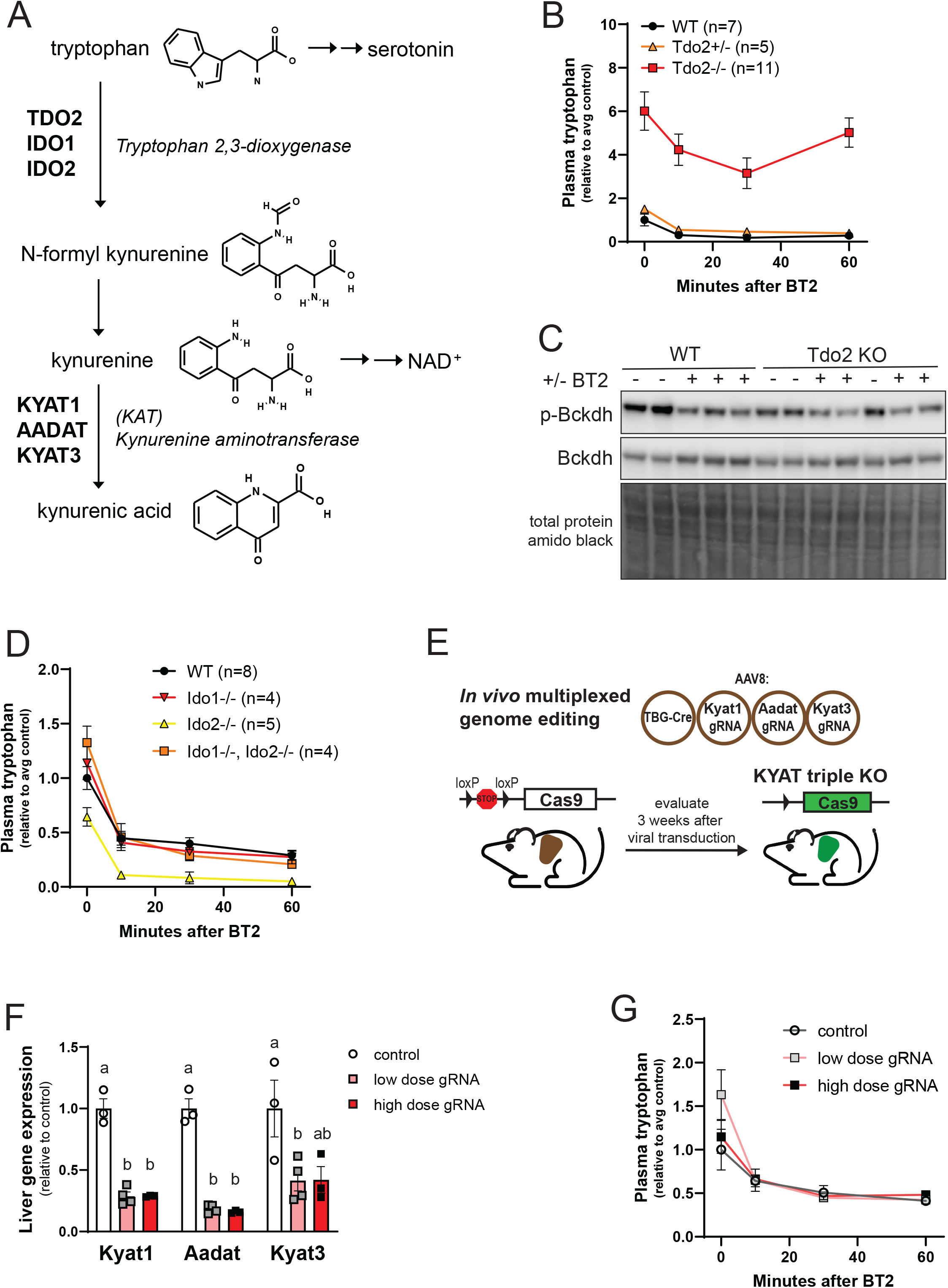
BT2 lowers plasma tryptophan independent of rate-limiting tryptophan catabolic enzymes. (**A**) Tryptophan catabolic pathway to kynurenic acid. (**B**) Acute BT2 treatment of Tdo2^-/-^ mice (n=5-11). (**C**) Immunoblot of liver protein 90 minutes after 40 mg/kg gavage of BT2 in Tdo2^-/-^ and littermate controls demonstrates TDO is not required for inhibition of BCKDH phosphorylation by BT2. (**D**) Acute BT2 treatment of Ido1^-/-^, Ido2^-/-^ and Ido1/2 double knockout mice lowered plasma tryptophan (n=4-8). (**E**) The kynurenine aminotransferases Kyat1, Aadat, and Kyat3 were simultaneously deleted in the liver (triple KO) by CRISPR/Cas9-mediated knockout. (**F**) Reduced liver mRNA abundance in triple KO animals. (**G**) Efficient BT2-induced decrease in plasma tryptophan in triple KO animals. Low dose = 1.5E12 genome copies (gc) and high dose = 3E12 gc; n= 3-4. Letters indicate statistically significant differences by one-way ANOVA and Tukey’s multiple comparisons test (ie. a and b are statistically significantly different with p<0.05).

We therefore next tested the role of IDO1 and IDO2 in BT2-mediated effects on tryptophan catabolism. *Ido1*^-/-^, *Ido2*^-/-^, and *Ido1*^-/-^;*Ido2*^-/-^ double knockout (DKO) mice (previously described ^40,41^), along with littermate controls, were treated with oral gavage of 40mg/kg BT2 followed by serial plasma sampling (Figure 2D). While tryptophan levels were lower at baseline in *Ido2*^-/-^ mice, the plasma tryptophan levels in mice of all three genotypes were reduced by BT2 treatment to a similar extent as in control mice (Figure 2D). Together, these results demonstrate that genetic ablation of any of the tryptophan 2,3-dioxgenase or indole 2,3-dioxygenase enzymes does not prevent the BT2-induced decrease in circulating tryptophan.

The increase of kynurenic acid levels in the plasma after BT2 treatment suggested that, alternatively, perhaps BT2 activates one or more of the kynurenine aminotransferases (Kyats), which mediate transamination of kynurenine to kynurenic acid in the liver ^35,42^. There are 4 main enzymes with kynurenine aminotransferase activity in mouse liver: Kyat1, Aadat, Kyat3, and Got2. We therefore proceeded to generate mice with liver-specific deletion of Kyat1, Aadat, and Kyat3. Got2 is the mitochondrial aspartate aminotransferase, with a wide range of substrates and pleiotropic effects, so we chose not to target that enzyme *in vivo*. We generated these mice using the recently described CRISPR/Cas9-mediated rapid *in vivo* multiplexed editing (RIME) system ^43^ (Figure 2E). Briefly, guide RNAs to knockout Kyat1, Aadat, and Kyat3 were generated and integrated into AAV8 vectors for liver-enriched expression. Mice with homozygous lox-stop-lox-Cas9 alleles in the ROSA26 locus were simultaneously infected with gRNA expression vectors and AAV8 encoding for Cre recombinase under control of the liver-specific TBG promoter which induced expression of Cas9 in liver. This approach resulted in efficient deletion of all three Kyats specifically in the liver by three weeks after induction (Figure 2F). Liver-specific deletion of the three Kyats did not affect plasma tryptophan levels at baseline, and, importantly, BT2 once again suppressed circulating tryptophan levels to a similar extent as control mice (Figure 2G). We conclude that BT2 promotes tryptophan catabolism by a mechanism other than direct activation of any principal rate-limiting tryptophan catabolic or kynurenine-consuming enzymes.

### BT2 binds to albumin and displaces tryptophan

Unique among amino acids in the circulation, 66-90% of tryptophan is bound to plasma proteins, specifically to site II on albumin ^44,45^. We noticed that the structure of BT2 has similarities to tryptophan (Figure 1A and 2A). It was previously determined that ∼99% of BT2 is bound to plasma proteins ^17^. Considering this, and that tryptophan depletion is very rapid (Figure 1), we therefore hypothesized that BT2 may compete with tryptophan binding to plasma protein. To test this notion, we performed equilibrium dialysis, with a membrane permeable to 8 kDa or smaller, between phosphate-buffered saline (PBS) and fetal bovine serum (serum) and in the presence of BT2 or vehicle (DMSO) (Figure 3A). Subsequent measurements of the ratio of a given metabolite in the serum versus PBS compartments reveals the extent to which this metabolite binds to serum protein. BT2 itself was tightly bound to serum protein and partitioned into the PBS compartment with increasing concentration of BT2 (Figure 3B). Similarly, tryptophan was ∼70% bound to serum proteins under these conditions (3C). Addition of BT2, however, gradually displaced tryptophan from serum protein, completely displacing tryptophan at a BT2 concentration of 150 µM and 300 µM, roughly similar to the endogenous concentration of tryptophan (Figure 3C). In contrast, BT2 did not impact fatty acid binding to serum protein (Figure 3D), while lactate did not bind to serum proteins at all (3E). Similar effects were seen when using mouse serum instead of FBS, from control or from *Tdo2*^-/-^ mice that received a vehicle or BT2 gavage (Figure 3F). We conclude that BT2 specifically displaces tryptophan from serum protein at physiological doses.

**Figure 3.**
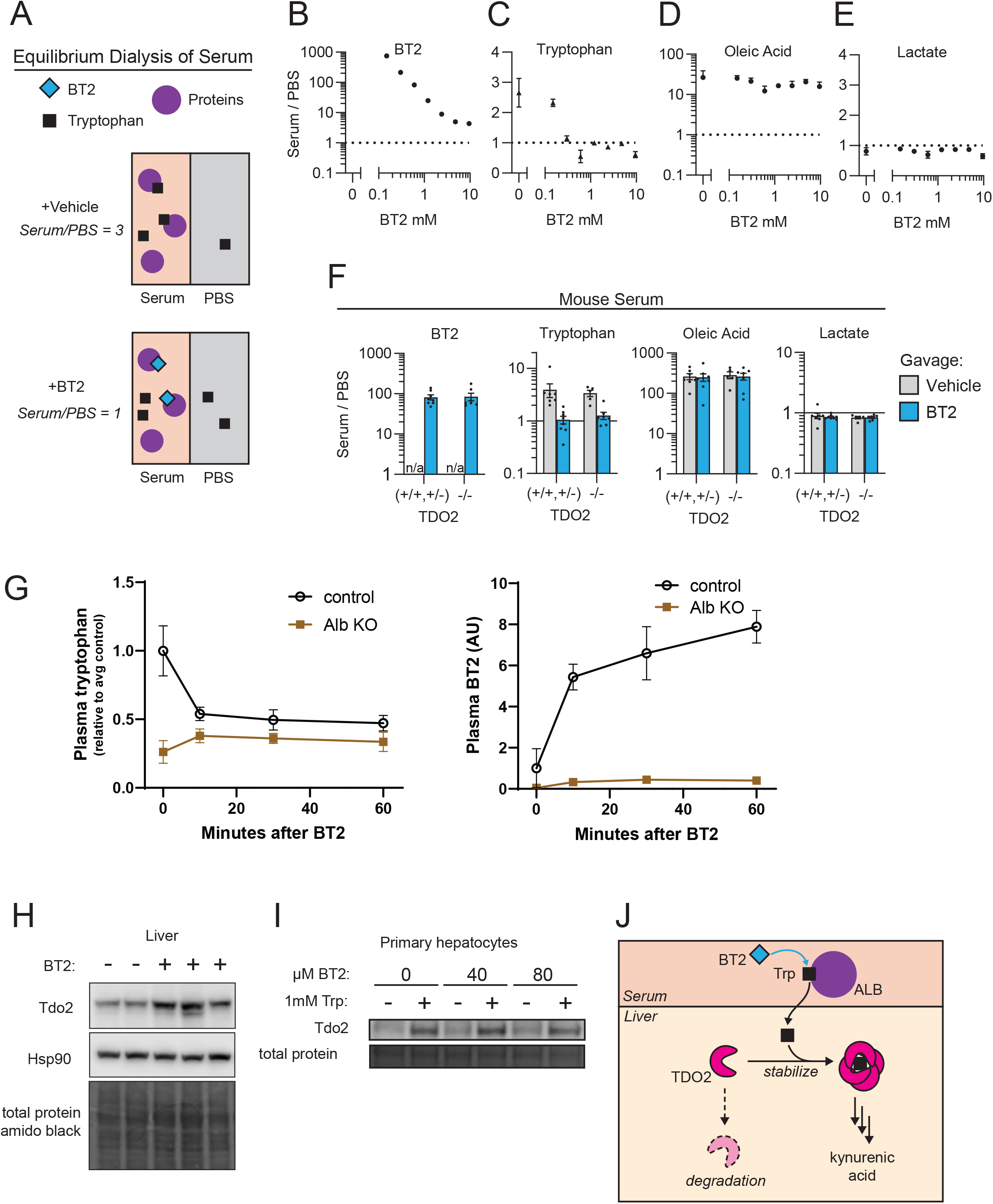
BT2 displaces tryptophan from serum albumin *in vivo*. (**A**) Equilibrium dialysis of serum can identify metabolites that BT2 displaces from serum proteins. After dialysis, the ratio of metabolite concentration in the serum compared to that in PBS indicates the retention of the metabolite in the serum chamber due to protein binding. (**B**) BT2 binds serum proteins avidly. Tryptophan binds serum proteins and is displaced by BT2 at 0.3 mM or higher concentrations. (**D**) Oleic acid binds to serum protein, and is not displaced by BT2. (**E**) Lactate does not bind to serum protein (n=3). (**F**) Equilibrium dialysis of serum collected 90 minutes after gavage of 40 mg/kg BT2 or vehicle from TDO2 KO and littermate control mice reveals BT2 displaces tryptophan but not oleic acid from serum protein. (**G**) BT2 is not retained in the plasma, and does not lower tryptophan levels, in albumin KO mice: mice were administered 40 mg/kg BT2 by oral gavage, followed by plasma tryptophan and BT2 measurements at the indicated times (n=5-6). (**H**) BT2 acutely stabilizes TDO2 protein *in vivo*. Immunoblot of mouse liver 90 minutes after oral gavage of BT2 (40 mg/kg). (**I**) Tryptophan increases Tdo2 protein expression in mouse primary hepatocytes, but BT2 does not. (**J**) Illustration of BT2 displacing tryptophan from albumin, followed by free tryptophan stabilization of TDO2 and subsequent clearance. In the absence of TDO2, tryptophan is likely cleared by IDO (not pictured).

To specifically probe the role of albumin binding *in vivo*, we next tested the effects of BT2 in mice lacking albumin. *Alb*^-/-^ mice are overtly phenotypically normal and have been used widely for pharmacology, immunology, and cancer biology research ^46-48^. Total plasma tryptophan levels in *Alb*^-/-^ mice were 25% of those in control mice (Figure 3G), consistent with the absence of the albumin-bound tryptophan pool. While treatment with BT2 suppressed tryptophan levels in control mice, BT2 administration had no effect on plasma tryptophan levels in *Alb*^-/-^ mice (Figure 3G), again consistent with the absence of an albumin-bound tryptophan pool to displace. Also consistent with the tight binding of BT2 to albumin, total plasma levels of BT2 achieved were much lower in *Alb*^-/-^ mice than in controls (Figure 3G). Together, these data indicate that BT2 displaces tryptophan from albumin *in vivo*, increasing the freely available form of tryptophan.

Free tryptophan is known to stabilize hepatic TDO2 protein ^49,50^, and indeed, we find that 90 minutes after BT2 gavage, liver Tdo2 protein levels were significantly increased (Figure 3H). In contrast, in cultured primary mouse hepatocytes, where tryptophan and BT2 delivery is not limited by albumin, BT2 did not increase protein levels of Tdo2, while free tryptophan did (Figure 3I). Together, these data strongly support the conclusion that BT2 displaces tryptophan from albumin in plasma, both *in vitro* and *in vivo*, thereby transiently increasing free tryptophan availability and promoting its catabolism by mass action^51^ (model: Figure 3J).

## Discussion

There is active interest from academic and industry groups in manipulating branched chain amino acid oxidation to treat several diseases, including insulin resistance, heart failure, hypertension, and cancer ^17,20,21,23,24,29,31,32,52,53^. Many of the data generated in these studies are based on experiments with BT2, under the usually implied assumption that it acts specifically on BCKDK. Our data now reveal that BT2 also displaces tryptophan from serum albumin, promoting its catabolism. Possible effects on tryptophan metabolism must therefore be considered when interpreting results of studies that use BT2. Testing with *Bckdk*^-/-^ mice can confirm if effects of BT2 are on-target. It will also be of interest to determine if newly developed BT2-like inhibitors of BCKDK also bind to albumin and displace tryptophan^31^. Other compounds are also known to bind the same site in albumin ^54^ and may also displace tryptophan. It should be noted, however, that the plasma concentrations of albumin and tryptophan are roughly 300 μM and 50 μM, respectively; relatively high doses of compounds are therefore needed to displace tryptophan, and drugs with high potency and low dose of administration are thus unlikely to meaningfully do so. Drugs that bind other sites on albumin, such as those bound by fatty acids, should also not affect tryptophan levels.

Free tryptophan is rapidly catabolized after release from albumin. Most of this released tryptophan is likely cleared by hepatic TDO2^35^, but some may also be cleared by IDOs in other tissues, as is implied by the observation that tryptophan is also partially cleared upon BT2 treatment in TDO2^-/-^ mice. In low tryptophan conditions, TDO2 in the liver is ubiquitinylated and degraded, while conversely tryptophan-mediated stabilization of TDO2 leads to rapid catabolism of tryptophan^49^. Why liver cells regulate tryptophan levels tightly is not clear, however. High tryptophan levels are well-tolerated, exemplified by the absence of adverse effects in a patient with loss-of-function mutation in TDO2 and more than 10-fold elevations in circulating tryptophan levels ^55^. Tryptophan dietary supplementation is generally safe ^56,57^. The immediate physiological consequences of elevated tryptophan levels upon administration of BT2 or other compounds may thus be minimal but require further study. The stabilization of iron- and heme-containing TDO2 may, for example, affect intracellular oxidative balance.

Interestingly, we find that *Bckdk*^-/-^ mice have elevated plasma tryptophan levels compared to controls (Figure 1D) ^32^, concomitant with their lower levels of BCAAs. Conversely, patients with maple syrup-urine disorder (MSUD), in whom BCAA catabolism is defective, have elevated BCAAs and low tryptophan (and other large neutral amino acids) ^58^. These data thus indicate crosstalk between BCAA and tryptophan metabolism. Competition by BCAAs and tryptophan at the L-type amino acid transporter (LAT1), known to occur at the blood brain barrier, is unlikely to be sufficient to explain these systemic observations.

In summary, we confirm that BT2 activates BCAA oxidation via inhibition of BCKDK, and we also report a robust activation of tryptophan catabolism by BT2, caused by the displacement of tryptophan from serum albumin, and independent of BCKDK or tryptophan catabolic enzymes. Studies with agents that bind serum albumin should consider the possible confounding effects of tryptophan displacement and catabolism.

## Materials and Methods

### Mice

All procedures were performed in accordance with the NIH’s *Guide for the Care and Use of Laboratory Animals* and under the approval of the Animal Care and Use Committees at the University of Pennsylvania and Rutgers University. Unless otherwise specified, wild-type (WT) C57BL/6J mice were obtained from the Jackson Laboratory (JAX# 000664), and 8-10-week old males were used for experiments.

### Generation of BCKDK KO mice

Two different lines of BCKDK knockout mice were used. For the experiment shown in Figure 1C, BCKDK knockout mice were from the line generated using gene-trap technology, originally described by Joshi et al^59^. For other experiments, the BCKDK knockout mice were from the line generated using the lox/cre system, originally described by Murashige et al.^60^

### Generation of Tdo2 KO mice

Tdo2 KO mice were generated using CRISPR/Cas9-targeted deletion of Tdo2 exons 1 and 2 on a C57BL/6J background by the CRISPR/Cas9 Mouse Targeting Core at the University of Pennsylvania. The 5’ and 3’ gRNA target sequences were as follows (target + PAM): CAGGCATGGCCCCTAGCTAAGGG and GAATTTGCAGATTTCCGATGGGG. Genotyping primers include forward primer CTACAGTCCTTGCCCTAACTTC and reverse primer AAGTCCTCCTTTGCTGGCTC with 1445 base pair (bp) amplicon in WT mice and 582 bp in Tdo2 KO. Presence or absence of the WT Tdo2 allele was also tested with forward primer GCCCCAACAAAAATCCACCC and reverse primer GTGTCTCAGGACCACACAGG for a product of 195 bp.

### Ido1, Ido2, and Ido1;Ido2 DKO mice

Ido1^-/-^ mice were obtained from the Jackson Laboratory (JAX# 005867). Ido2^-/-^ and Ido1^-/-^Ido2^-/-^ double knockout (DKO) mice were generously provided by Dr. Merlo and Dr. Mandik-Nayak^40,41^ and originally made by Dr. Peter Murray^61^.

### Generation of liver-specific triple-kynurenine aminotransferase KO mice

CRISPR/Cas9-mediated rapid *in vivo* multiplexed editing (RIME) ^43^ was used to generate mice with liver-specific deletion of kynurenine aminotransferase enzymes. Guide RNAs (gRNAs) to knockout Kyat1, Aadat, and Kyat3 were generated, tested in cultured mouse cells, and integrated into AAV8 vectors for liver-enriched expression. AAV8 was harvested and purified as described in ^43^. The sequences for the gRNAs were Kyat1:GAGTATGATGTCGTGAACTTGGG, Aadat:CAAGTCGTCTTCAAACCGGATGG, and Kyat3:AAAAACGCCAAACGAATCGAAGG. Mice with homozygous lox-stop-lox-Cas9 alleles in the ROSA26 locus (JAX # 026175) were simultaneously infected with AAV8-gRNAs and AAV8 TBG-Cre (1.5 or 3 x 10^12^ genome copies) by retroorbital injection to induce expression of Cas9 in liver and genome editing at the Kyat1, Aadat, and Kyat3 loci. Three weeks after injection, mice were administered BT2 by oral gavage, and plasma tryptophan was measured. After several days of recovery, mice were euthanized and tissues collected for validation of loss of function of the kynurenine aminotransferases. For qRT-PCR, total RNA was isolated using Trizol reagent (Invitrogen 15596026), cDNA synthesized with random primers (Applied Biosystems High-Capacity cDNA reverse transcription kit, 4368814), and 384-well reactions (total volume 5 µL) were prepared with 2X SYBR green master mix and gene-specific primers shown below. Data was normalized to the average C_t_ value for 18S and Rpl22 and expressed as 2^-delta C_t_ relative to control mice.

### Primers for qRT-PCR

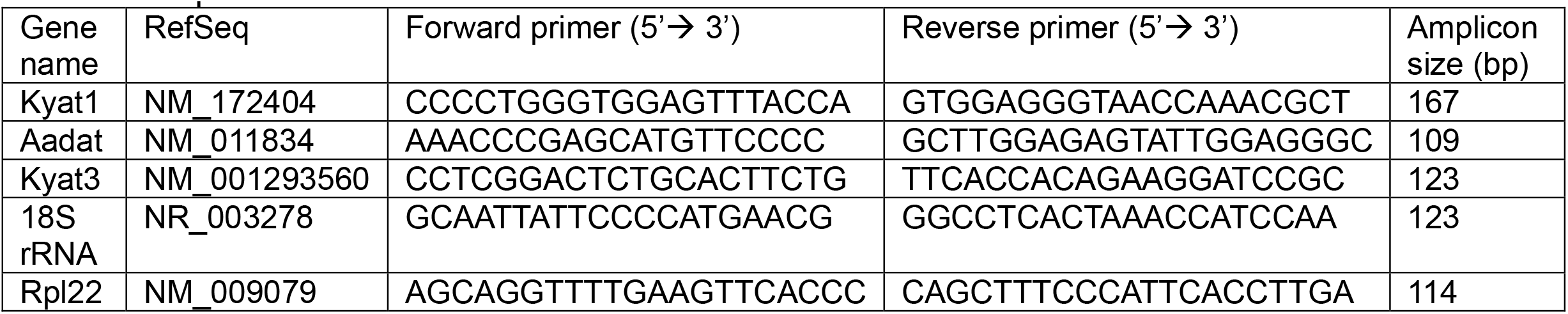

### Albumin KO mice

Albumin KO mice were obtained from the Jackson Laboratory (JAX# 025200), and males 8-15-weeks-old were used for experiments ^46-48^.

### Oral Gavage of BT2

BT2 (Chem Impex 25643) was dissolved at 200 mg/mL in DMSO, then diluted (5% volume per volume (v/v)) to a final concentration of 10 mg/mL in a suspension of 10% (v/v) Cremophor EL (Millipore 238470) and 85% (v/v) 0.1 M sodium bicarbonate, pH 9 (Gibco 25080-094). Mice were *ad lib* fed and given an oral gavage of BT2 at 40 mg/kg gavage around 13:00. Tail blood was collected just before gavage (0 min) and at the times indicated after gavage.

### Intraperitoneal Injection of BT2 or Sodium Phenylbutyrate (NaPB)

For intraperitoneal injection, BT2 (Sigma 592862), was prepared in sterile saline at 4 mg/mL (the BT2 purchased from Sigma is somewhat soluble in saline, but Chem Impex 25643 is less soluble) then 10 ul per gram body weight was injected intraperitoneally. Sodium phenylbutyrate (NaPB) (Sigma SML0309-100mg) was prepared in sterile saline at 12.5 mg/mL and a dose of 200 mg/kg was injected intraperitoneally (volume = mouse body weight x 16).

### Intravenous Injection of BT2

Mice were catheterized in their jugular vein and carotid artery one week prior to the experiment. On the day of the experiment, BT2 (Chem Impex 25643) was prepared as described for oral gavage of BT2 to a final concentration of 10 mg/mL. Mice were fasted 4 hours prior to the experiment, then a baseline sample of arterial blood was collected. BT2 (40 mg/kg) was slowly injected by hand through the catheter in the jugular vein. Additional samples of arterial blood were collected at the indicated times.

### Plasma preparation for LC or GC-MS

Tail blood was collected in lithium-heparinized capillary blood collection tubes (Sarstedt #16.443.100) and stored on ice. Samples were spun at 10,000 x g for 10 min at 4 °C. Plasma was transferred to another tube and analyzed immediately or stored at -80°C before sample preparation.

### LC-MS methods

Plasma or Dialysate levels of BT2, tryptophan, and other small metabolites were measured from 5 uL samples extracted with 200 uL of dry-ice cold analytical-grade methanol. Samples were incubated on dry ice for 15 minutes, then centrifuged at 21,000 x g for 15 minutes at 4°C. The supernatant was collected for LC-MS analysis. Samples from dialysis experiments were additionally diluted with 1× methanol prior to analysis.

Metabolites were measured on an orbitrap mass spectrometer (Q Exactive Plus, Exploris 240, or Exploris 480) coupled to a Vanquish UHPLC system (Thermo Fisher Scientific) with electrospray ionization and scan range m/z between 60-1000 at 1 Hz, with at least 140,000 resolution. Data was generated from both positive and negative ion mode. LC separation was performed on an XBridge BEH Amide column (2.1×150mm, 2.5 um particle size, 130 Angstrom pore size; Waters Corporation) using a gradient with solvent A (95:5 water:acetonitrile with 20 mM ammonium acetate and 20 mM ammonium hydroxide, pH 9.45) and solvent B (acetonitrile). Flow rate was 150 ul/min. A typical LC gradient was: 0 min, 85% B; 2 min, 85% B; 3 min, 80% B; 5 min, 80% B; 6 min, 75% B; 7 min, 75% B; 8 min, 70% B; 9 min, 70% B; 10 min, 50% B; 12 min, 50% B; 13 min, 25% B; 16 min, 25% B; 18 min, 0% B; 23 min, 0% B; 24 min, 85% B; and 30 min, 85% B. Injection volume was 5-10 uL and autosampler was kept at 4C. Data was analyzed using El-Maven (version 12).

### GC-MS detection of tryptophan and BT2

Metabolites were extracted from 10 µL of mouse plasma with 10.3 uL of 0.12 N hydrochloric acid followed by addition of 120 µL of dry-ice cold analytical-grade methanol. Each sample was spiked with 5.6 nmol of L-norvaline as an internal standard, provided in the methanol. Samples were incubated on dry ice for 15 min then centrifuged for at 21,000 x g for 15 min at 4°C. Supernatants containing soluble metabolites were transferred to new tubes and dried under vacuum in a speedvac microcentrifuge concentrator. In a fume hood, dried samples were resuspended in 40 µL of room-temperature 1:1 (v/v) analytical-grade acetonitrile and MtBSTFA (N-methyl-N-(tert-butyldimetylsilyl)trifluoroacetamide) (Regis Technologies 1-270242-200), and were heated on a 70°C heat block for 90 min. Then samples were cooled to room temperature (about 5 min), centrifuged at 13,000 x g, and the supernatant was transferred to GC-MS vials with polypropylene inserts for small volume samples.

One microliter of the sample was injected via automatic liquid sampler (Agilent 7693A) into an Agilent 7890B gas chromatograph (GC) coupled with an Agilent 5977B mass selective detector (MSD) (Agilent Technologies). The GC was operated in splitless injection mode with helium as the carrier gas at a flow rate of 1.2mL/min. The GC column was a 30 m x 250 µm x 0.25 µm HP-5ms Ultra Inert column (Agilent 19091S-433UI). The inlet temperature was 250°C, and, after 3 min at 100°C, the oven temperature program was increased as follows: 20°C/min to 210°C then 4°C/min to 300°C and hold 5 min. The transfer line temperature was 250°C, and the MSD source and quadrupole temperatures were 230°C and 150°C, respectively. After a 6 min solvent delay, the MSD was operated in electron ionization mode and scan mode with a mass range of 50-550 AMU at 2.9 scans/s. Agilent MassHunter Qualitative Analysis software (B.07.00) was used for visualization of chromatograms. BT2 derivatized with MTBSTFA has a characteristic m/z of 303 and 305 (doublet for ^35^Cl and ^37^Cl) at 18.3 minutes under these conditions.

### Equilibrium dialysis

Equilibrium dialysis was performed using Rapid Equilibrium Dialysis kits (ThermoFisher Scientific 90006) according to manufacturer instructions. For the dialysis of fetal bovine serum (FBS), BT2 (from working stock 200 mg/mL DMSO) or vehicle was spiked into aliquots of FBS and vortexed. 100 uL samples were dialyzed with 350 uL of phosphate-buffered saline (samples added to red chambers, saline added to white chambers). For the dialysis of mouse serum, 50 uL of serum was dialyzed with 300 uL of phosphate-buffered saline. All samples were dialyzed with gentle shaking at 37 degrees C, protected from light, for four hours. 40 uL aliquots were then collected from every chamber and frozen immediately at -80 degrees C until they were processed for LC-MS analysis.

### Primary hepatocyte isolation and culture

Mouse primary hepatocytes were isolated and cultured as previously reported ^62^. Tryptophan and BT2 were administered at the concentrations shown for 24 hours using water and methanol as vehicle controls, respectively.

### Western blot

Frozen liver tissue was homogenized in RIPA buffer (RPI R26200) containing cOmplete Mini, EDTA-free protease inhibitor (Roche, 11836170001) and PhosSTOP phosphatase inhibitor (Roche, 4906837001) with ceramic beads in 2 mL screw-cap tubes using the Omni Bead Ruptor 12 for two 20-second rounds of homogenization. The homogenate was centrifuged at 14,000 x g for 15 min at 4 ºC. The supernatant (lysate) was transferred to a new tube, and protein concentration was determined by BCA protein assay kit (Thermo Fisher, 23225). Cells were lysed in the same RIPA buffer with protease and phosphatase inhibitors, sonicated briefly, centrifuged to collect supernatant for quantification and immunoblotting.

For immunoblots, 15-25 µg of total protein from mouse tissue or cell lysates was loaded in 4-20% polyacrylamide gradient Bis-Tris gels and electrophoresed at 130V for 1 to 1.5 hours. The gels were transferred to PVDF membranes (Immobilon-P, IPVH00010) at 100V for 90 minutes, which were then blocked in 5% non-fat dry milk in TBST for 60 minutes and incubated with primary antibody (1:1000) overnight at 4 ºC. The primary antibodies used include Tdo2 (Proteintech 15880-1-AP), Hsp90 (Cell Signaling Technology 4874), phospho-Bckdh (BCKDE1A, Ser293) (Bethyl A304-672A), and total Bckdh (BCKDE1A) (Bethyl A303-790A). After overnight incubation with primary antibody, membranes were washed in TBST and incubated for 60 minutes with rabbit horseradish peroxidase (HRP)-conjugated secondary antibody (Cell Signaling Technology 7074) diluted at 1:2000 in 5% milk-TBST. The membranes were washed again in TBST, incubated for ∼30 seconds with enhanced chemiluminescent substrate for HRP (Thermo Fisher, 34094 or 34075), and imaged using a digital imager. For amido black total protein staining, PVDF membranes were incubated in 0.1% naphthol blue black (w/v), 10% methanol (v/v), 2% acetic acid (v/v) for 5 min and destained in 10% acetic acid (v/v), 25% isopropanol (v/v) for 1 min before imaging.

### Statistical analysis

Graphs were generated in GraphPad Prism, and mean ± SEM is displayed unless indicated otherwise. Oral gavage of BT2 experiments in which there are both time and genotype factors and repeated measures were analyzed a mixed-effects model with Geisser-Greenhouse correction. P-values are reported for statistically significant genotype effects.

## Author Contributions

Conceptualization: CEB, MN, ZA; Investigation: CEB, MDN, CJ, JP, MCB, ETM, WOJ, QC; Methodology: CEB, MDN, QC; Resources: LM, LMN, TGA, JDR, ZA; Writing—original draft: CEB, MDN, ZA; Writing—review and editing: all authors.

## Acknowledgements

We thank the Penn CRISPR Core for generating the Tdo2 KO mice and researchers from Laura Mandik-Nayak’s lab at Lankenau Institute for Medical Research for assistance with the Ido2 KO and Ido1; Ido2 DKO mice. We thank Takeshi Katsuda of Ben Stanger’s lab at University of Pennsylvania for guidance with CRISPR/Cas9-mediated rapid *in vivo* multiplexed editing (RIME) and for providing plasmids and mice, and we thank Sarmistha Mukherjee of Joseph Baur’s lab at University of Pennsylvania for expertise on primary hepatocyte culture. We thank Lingfan Liang for guidance in dialysis experiments. We also thank members of the Arany lab for helpful discussion and technical assistance with this project, especially Jian Li, Senali Dansou, and Michael Noji.

